# Tagging Very Small Fish: Two Effective and Low Impact Methods

**DOI:** 10.1101/437772

**Authors:** Deijah D. Bradley, Elianna J. Schimke, Alyssa P. Alvey, Hans A. Hofmann, Tessa K. Solomon-Lane

## Abstract

Identifying individuals over time and across contexts is essential in many scientific fields. There are a variety of well-established methods for uniquely marking individuals (e.g., visible implant elastomer, barcodes, paint). However, for some species, life history stages, and/or experiments, existing methods are not sufficient. Here, we describe procedures for how two tagging methods – a tattoo ink injection method and a fishing line piercing method – can be used with the youngest, smallest juveniles of the African cichlid fish, *Astatotilapia burtoni*, which are too small for the methods used with adults. With the tattoo method, we injected tattoo ink into the dorsal muscle. Different colors and injection locations can be used to distinguish among individuals over a period of weeks (up to 4 weeks, average 2.5-3 weeks under our conditions). Because fish this young and small are sensitive to handling and injection, we also include physiological data showing fish recover well from anesthetization and tagging. With the piercing method, very thin fishing line is threaded through the dorsal muscle and tied into a barbell or loop. Unique colors and patterns can be used to distinguish among individuals over a period of months. Because a physical tag might impede normal movement in a very small fish, we also include data from an open field exploration test showing similar behavior between tagged and control (non-tagged) juveniles. We expect these effective and inexpensive methods to be useful for a variety of small species and will facilitate early-life, developmental, and longitudinal research.

## INTRODUCTION

The ability to uniquely identify individuals is essential to many scientific endeavors. In some species, the natural, external patterns on the body are sufficiently distinctive for identification, such as humpback whales, *Megaptera novaeangliae* (Katona et al., 1979), the African cichlid fish, *Neolamprologus pulcher* (Kohda et al., 2015; Balzarini et al., 2017), the guppy, *Poecilia reticulata* (Kemp et al., 2009), and the bluebanded goby, *Lythrypnus dalli* (Reavis and Grober, 1999). For most species, however, it is necessary to mark individuals to reliably identify them over time and across contexts. The ideal tag allows for easy and unambiguous identification, lasts the duration of the experiment(s), and interferes minimally with the animal and experimental conditions (Malone et al., 1999). The method also must be ethical and in compliance with the local regulations for the care and use of experimental animals. There are a variety of useful and well-established methods for marking individuals. For example, paint or dye is used in diverse species for external marking, from insects to mammals (e.g., Jones et al., 1996; Williamson et al., 2016); combinations of colored and metal bands are placed on bird legs (Frazier, 2015); fish fins can be uniquely clipped (e.g., Hammer and Lee Blankenship, 2001; Thompson et al., 2005); visible implant elastomer tags can be injected under the skin or scales of amphibians, reptiles, and fish (e.g., Malone et al., 1999; Thompson et al., 2005; Grant, 2008); unique identifiers (e.g., numbers, barcodes) can be glued to the animal (e.g., Formica et al., 2012); passive integrated transponders (PIT) can be implanted and detected by radio signal (e.g., Jørgensen et al., 2017; Kraus et al., 2017; Lennox et al., 2025); surgical steel coded wire implant tags (e.g., Martin and Wainwright, 2013; Martin and Gould, 2020), and more (e.g., Hasler and Faber, 1941; Volk et al., 1999). Developments in automated tracking technology have also influenced identification methods, such as radio frequency identification (RFID) tags (e.g., Lewejohann et al., 2009; Weissbrod et al., 2013) and computer-vision systems (e.g., bleach-marks on mouse fur, Ohayon et al., 2013). The spatial and temporal scales involved, and the requirements and limitations of the study species and experiment, together determine the most appropriate marking method.

Despite the variety of existing options, there are barriers for using these methods in some species, life history stages, and/or experiments. Here, we describe two procedures – a tattoo method and a piercing method – that make it feasible to tag even the youngest, smallest juveniles (average size: 8.31 ± 0.039 mm standard length; 0.022 ± 0.0005 g mass, Fig 1A) of the African cichlid fish, Burton’s Mouthbrooder (*Astatotilapia burtoni*), a model system in social neuroscience (Hofmann, 2003; Maruska and Fernald, 2018). Adults of this species are routinely tagged using visible implant elastomer tags, a colorful liquid that dries as a pliable solid (Northwest Marine Technology, Inc.), or a colored bead secured with a plastic tag through the dorsal muscle (as in Trainor and Hofmann, 2006). However, very small juvenile *A. burtoni* do not have sufficient dorsal muscle into which to inject elastomer (Fig 1B-E), which uses a 30-gauge needle because of the viscosity of the substance. The bead (often cut in half to be lighter) and plastic tag (can also be used alone) are too large and heavy for the juveniles.

**Figure 1:**
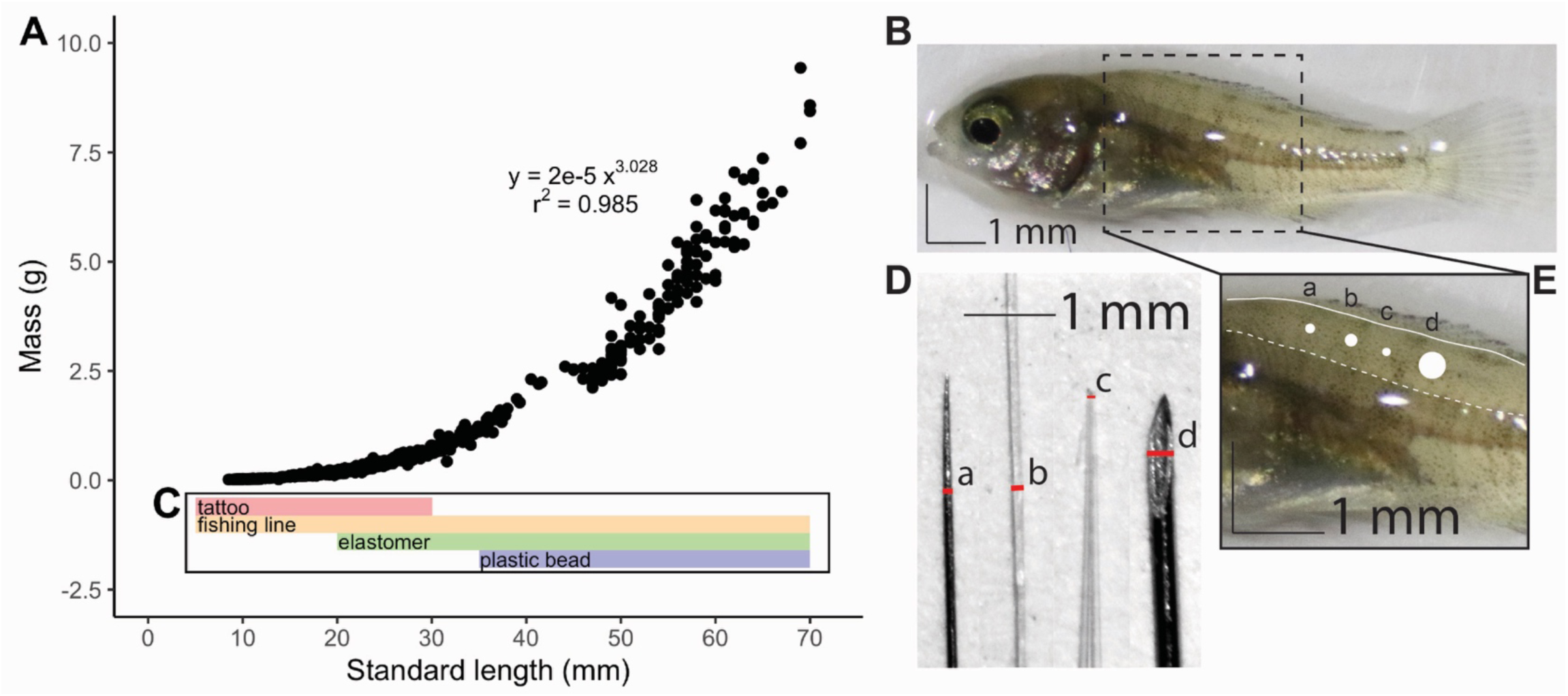
Summary of appropriate tagging methods for *Astatotilapia burtoni* of different body sizes. A) Association between standard length (mm) and mass (g) (n=729). B) Image of 1-day old juvenile (8.8 mm standard length). C) Four methods used for tagging *A. burtoni* at different body sizes and life history stages, including the two methods described here: tattooing and a fishing line piercing through the dorsal muscle. The size upper ranges for the tagging methods are approximate and may vary depending on the experiment (e.g., short- vs. long-term, live vs. video behavior scoring) and the species. D) Image of a 0.1 mm diameter minutien pin (a: for piercing the smallest juveniles), fishing line (b: for tagging the smallest juveniles), pulled glass capillary tube needle (c: for tattooing), and a 31-gauge needle (d: for piercing larger juveniles). The red lines indicating diameter are the same diameter as the white circles in (E) (D and E are at the same scale). E) Cropped image of the fish in (B). The dorsal edge of the spinal cord is edged in dashed white lines. The dorsal edge of the fish is shown in the sold white line. White circles show the diameter of the minutien pin (a), fishing line (b), capillary tube needle (c), and 31-g needle (d) shown in (D).

The tattoo method adapts existing approaches that use injected or implanted dyes or paints for identification (Lotrich and Meredith, 1974; Ryan, 1975; Hill and Grossman, 1987; Chart and Bergersen, 1988) for use on juvenile fish that are at least an order of magnitude smaller than the animals used in previous studies. We describe how to inject nanoliter quantities (∼100 nL) of tattoo ink into the dorsal muscle using a pulled glass capillary tube needle. We can use this method with the youngest and smallest fish, including larva that are still developing. This method is moderately long-lasting: up to 4 weeks, with an average of 2.5-3 weeks under our laboratory conditions. This duration is similar to injection methods in other species (Thresher and Gronell, 1978). This tag does not appear to interfere with locomotion, and it is visible to the naked eye. Because fish of this early developmental stage are especially sensitive to handling and prone to injury or death, we also present physiological data on how anesthetized and injected juveniles recover from the procedure compared to anesthetized juveniles that were not injected (Sneddon, 2012). All injected and control fish are expected to recover fully from anesthesia, and we hypothesize that tattoo injections would not significantly affect the timing of the stages of recovery from anesthesia.

The piercing method adapts existing approaches that use a physical tag for identification (e.g., Martin and Wainwright, 2013; Martin and Gould, 2020) for use in juvenile fish that are at least an order of magnitude smaller than the animals used in previous studies. Here, we describe how to thread very thin fishing line through the dorsal muscle of the fish using a minutien pin or needle appropriate to the fish’s size. The line is then tied into a barbell or loop and remains in place. These tags are visible to the naked eye, visible on camera for video analysis (depending on the size of the tag and cameras), it can remain in place as the juvenile grows, and it is sufficiently long-lasting for the duration of our experiments (months). Because a physical tag might impede locomotion in the smallest fish, we also include data from an open field exploration test using fully developed juveniles, 1-day after their removal from the mother’s buccal cavity. We chose fish this young because they are the most likely to be physically impeded by a tag and/or negatively affected by the procedure. We hypothesized that tagged juveniles would behave similarly to untagged fish in the open field, spending similar amounts of time in the different zones of the tank. Anesthesia recovery data are not included for this method.

Overall, we expect that these procedures will be useful to other researchers working with a variety of species and that they will continue to be developed, adapted, and improved. The ability to identify individuals over time, starting in early life, will make it more feasible to carry out developmental and longitudinal research when working with very small, young fish.

## METHODS

### Astatotilapia burtoni juveniles

We used *A. burtoni* juveniles from a laboratory population descended from a wild-caught stock (see lab strain from Hofmann and Renn: Pauquet et al., 2018). The adults that bred these juveniles were housed in naturalistic social groups of males and females. The mother orally incubates the larvae as they develop for 10-13 days after fertilization (Fernald and Hirata, 1979; Renn et al., 2009). When the larvae are fully developed and ready to leave the mother’s buccal cavity, the average standard length (SL) in our laboratory populations is 8.31 ± 0.039 mm (n=356, max SL: 10.18 mm, min SL: 4.6 mm, Fig 1A). All procedures were done in compliance with the Institutional Animal Care and Use Committee (IACUC) of the Department of Natural Sciences of Pitzer and Scripps Colleges (protocol #19-001) and the IACUC of The University of Texas at Austin.

### Tagging by tattoo ink injection

#### Tattoo ink and Nanoject injection set-up

We developed this procedure using commercially available sets of tattoo ink, including seven color sets of Millennium Mom’s Tattoo Ink (Amazon.com, $40 USD at the time of the work) and HAWINK Tattoo Ink (Amazon.com, $20 USD at the time of the work). These sets included vibrant colors such as red, yellow, green, and blue (Supplemental Fig 1), as well as neutral colors such as white, gray, and black. We also used Keep It Wet (Eternal Tattoo Supply, $5 USD at the time of the work) to keep the ink from drying as quickly, without affecting the color.

Tattoo injections were performed under a dissection microscope using the Nanoject II Auto-Nanoliter Injector (Drummond Scientific Company). The Nanoject was positioned on a micromanipulator attached to a magnetic base, which was secured to the benchtop on a magnetic board. See Supplemental Materials for detailed set-up information. Briefly, we used a puller (World Precision Instruments PUL-1) to make glass capillary tube needles (1.2 mm x 68 mm, A-M Systems). The tip of the needle was broken off using forceps, and then we backfilled the needle with mineral oil and secured it onto the Nanoject (Fig. 2A, Supplemental Fig 2). After emptying a portion of the mineral oil, we filled the needle with tattoo ink from a weigh boat or Parafilm. We found that different ink colors and brands had different consistencies and drying times in the needle, so we used a ratio ranging from 1:1 to 1:4 ink drops to water drops (or Keep It Wet additive). We performed multiple test injections (Supplemental Fig 2F) at the start and between fish to ensure the ink was not dried, clogging the needle.

**Figure 2:**
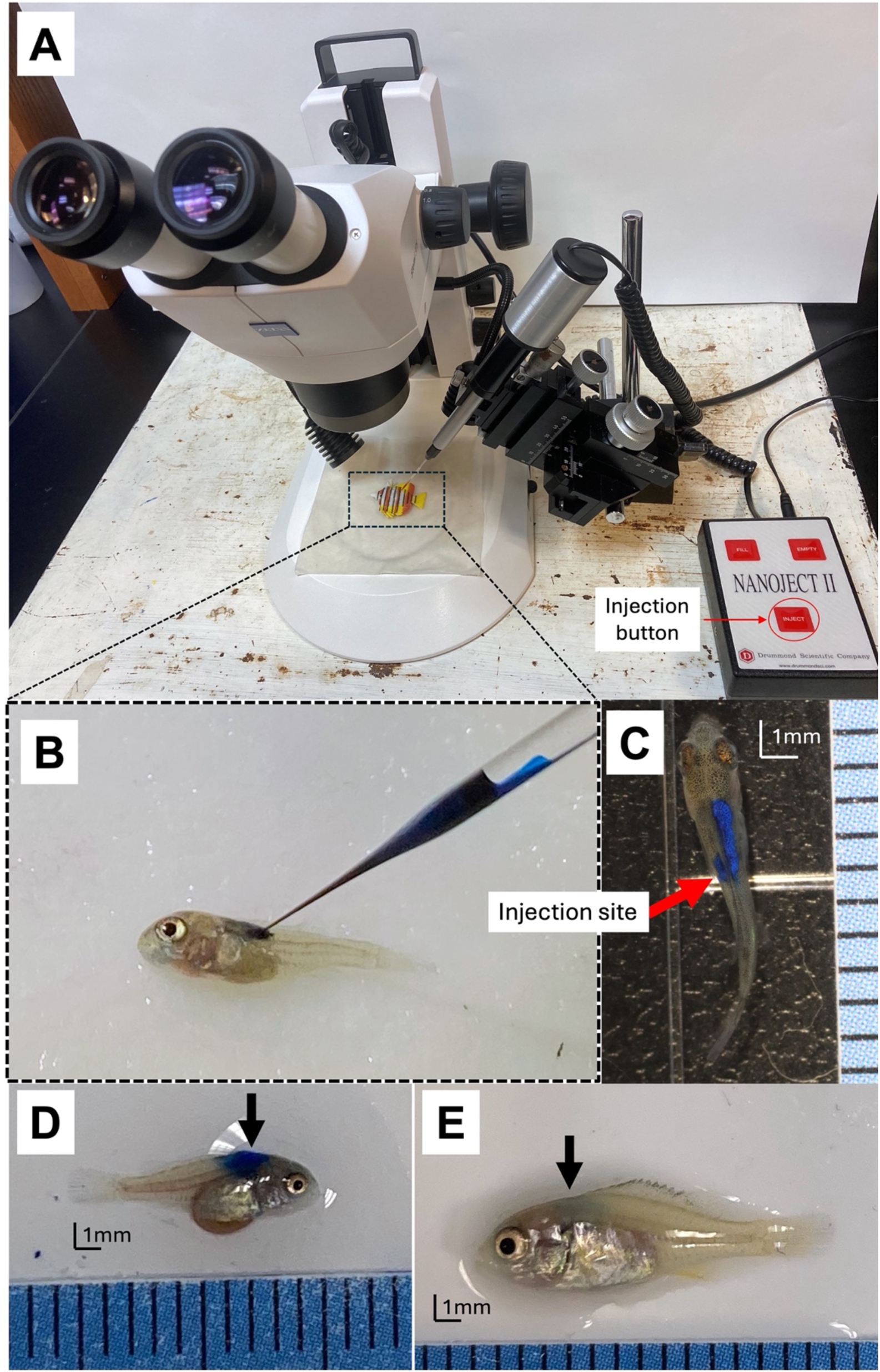
Tattoo ink injection tagging. **A)** Image of the Nanoject setup under a dissection scope, with a (plastic) fish placed on a wet napkin on the platform, prepped for injection. The injection button is labeled on the Nanoject controller. **B)** A juvenile fish being injected with blue ink using a pulled glass capillary tube needle. **C)** Fish immediately after injection with tattoo ink. The injection site is shown on the left, with the dye spreading on the right side of the fish. **D)** A juvenile fish 2 days after injection. **E)** The same juvenile fish as in (D, at the same scale) 36 days after injection. See Supplemental Information for pictures of additional tag colors.

#### Injection of tattoo ink

We removed the fish from the anesthetic (see below) and placed it on the injection platform under the dissection scope on a paper towel wet with aquarium water (Fig 2A). We positioned the fish so the needle could be inserted (using the micromanipulator) between scales and into the dorsal muscle, at the desired location along the rostral-caudal axis (Fig 2B). Young *A. burtoni* juveniles are semi-transparent at this developmental stage; therefore, the needle position was also checked visually prior to injection (Fig 2B). We then injected ∼100 nL of ink with 2 sequential injections of 50.6 nL (the largest volume for the Nanoject II model). The ‘tattoo’ was immediately visible at the injection site, as well as on the opposite side of the fish (Fig 2C-E). We kept the needle in place for ∼5 s to help prevent the ink from flowing out of the injection site. We then carefully placed the fish into a beaker of fresh aquarium water and aided recovery from anesthesia using artificial ventilation (see below). We wiped the needle with ethanol on a Kimwipe (Kimberly-Clark) between injections and changed the needle every 4-8 animals, as the needle dulled, if the needle clogged, or to change ink color.

#### Anesthesia and post-tattoo ink injection recovery

We anesthetized juveniles less than two weeks old in tricaine methanesulfonate (MS-222, Sigma Aldrich) at a dose of 0.0003 g / mL aquarium water, buffered with sodium hydroxide to pH 7-7.5. During anesthetization, we recorded the time it took for body movement to stop; loss of equilibrium, when the fish’s dorsal side first rolled away from an upright orientation; and when ventilation stopped, indicated by movement of the opercula ceasing fully. We removed fish from the MS-222 immediately after ventilation stopped.

After tagging (see above), the fish was placed in a 200 mL plastic beaker of fresh aquarium water to recover. For experiments that require strict procedural standardization, we recommend keeping the amount of time out of water consistent across fish (e.g., 1 min for tattooing only; 2 min for tattooing and an additional procedure, such as drug administration). Otherwise, we placed the fish back in aquarium water for recovery as quickly as possible.

We immediately began artificial ventilation using a plastic pipette to gently push water over the gills. Recovery from MS-222 is a highly stereotyped process (McFarland, 1959). We recorded the time it took for the fish to first move its body; first move the opercula; ventilate independently without the artificial ventilation of the pipette; initial regaining of equilibrium, when the dorsal side of the fish first returned to an upright position; and sustained equilibrium, when the fish was permanently upright. We also recorded when the fish looked fully recovered, which tended to happen suddenly. Fully recovered fish swam smoothly and with steady ventilation and no wobbles in equilibrium.

### Tagging by piercing with fishing line

#### Making the fishing line tag

Unique tags can be made from fishing line by coloring or dying the line, or purchasing fishing line manufactured in different colors (options may be limited). We used permanent markers (Sharpie) to uniquely color white fishing line, although this color fades over a period of weeks (3-6). “Beads” of different colors, sizes, or patterns can also be made along the length of the line by tying a knot (or more than one) and using a drop of super glue to preserve the color of the permanent marker. Non-toxic nail polish can also be used over knot(s) in the line for additional colors and patterns of “beads,” which are larger and more visible on camera than knots in the line alone.

For the juveniles that are too small for a 31-gauge insulin needle (or the smallest commercially available needle) to pass through the dorsal muscle (Fig 1D, E), we used a minutien pin (Fine Science Tools, 0.1mm diameter, 500 pins for $32 USD at the time of the work) in combination with the thinnest available fishing line (Berkley NanoFil Uni-filament fishing line, 2 lb break weight, 0.051 mm average diameter, 150 yards for $20 USD at the time of the work). For this method, the fishing line and pin are not physically attached.

For *A. burtoni* juveniles that are large enough for a 31-gauge insulin needle to pass through the dorsal muscle (body size > ∼12 mm SL, ∼0.08 g mass for *A. burtoni*), we attached the fishing line to a needle, making it easier and more efficient to tag many fish in a row. This design is modeled on an eyeless suture, which can be expensive to purchase, but our alternative is easy and inexpensive to make. We used Berkley NanoFil Uni-filament Fishing Line (4lb break weight, 0.1 mm average diameter, in the color clear mist; 150 yards for $20 USD at the time of the work) and BD Ultra-Fine^TM^ Short Needle (8 mm, 31G) Insulin Syringes (3/10 mL) (90 count for $21 USD at the time of the work). Any fishing line and needle combination can be used, as long as the average diameter of the line is smaller than the inner diameter of the needle.

To attach the fishing line to the needle, we removed the plunger from the syringe and threaded the line through the sharp end of the needle under a dissection scope. Line that was freshly and smoothly cut with a new razor blade was the easiest to work with. If the fishing line did not easily fit into the needle, we discarded a length of line and started at a new place along the spool. The diameter of the line varies slightly throughout, and it was easiest to work at an average or narrower section. Once started, we kept threading the line through the needle until it was visible in the barrel of the syringe (Supplemental Fig 3A). We then used forceps to grip the base of the needle, being careful not to dislodge the fishing line. We rocked the needle back and forth with the forceps until it detached from the syringe (Supplemental Fig 3B). From the blunt side of the needle, we then pulled the line through until it was sufficiently long (30-50 cm of line can be used to tag ∼4-10 fish, depending on the efficiency of the tagger). Finally, we cut the line from the spool using a razor blade and pulled the last bit of line through the needle until it was no longer sticking out beyond the sharp end. We did not find it necessary to crimp the blunt end of the needle to keep the line in place.

#### Anesthesia and recovery

We anesthetized juveniles less than 2 weeks old as described above. Older, larger juveniles were anesthetized in MS-222 at a dose of 0.0006 g / mL aquarium water, buffered to pH 7-7.5. We removed the larger juveniles from the MS-222 immediately after they stopped responding to touch, which occurred after losing equilibrium and after gill ventilation stopped. We aided fish in their recovery from anesthesia using artificial ventilation, as described above.

**Figure 3:**
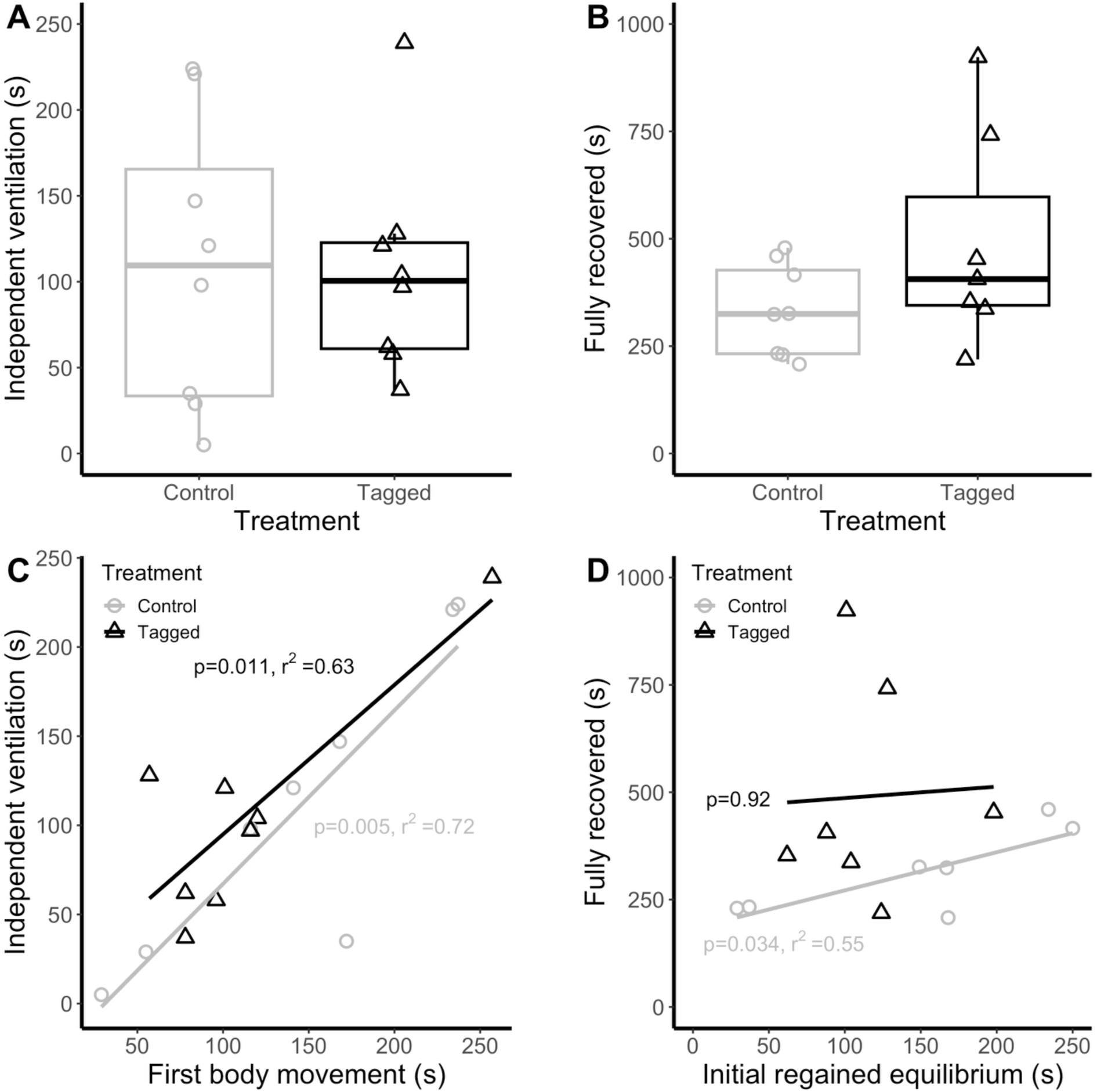
Recovery from anesthesia following tattoo ink injections: Time until A) ventilating independently and B) full recovery from anesthetization in control and tagged juveniles. C) Association between initial body movement and independent ventilation. D) Association between initial regained equilibrium and full recovery. All recovery times start when the fish was placed in the recovery beaker, following anesthetization and 1 min out of water. Tagged fish were injected with dye during the 1 min out of water.

#### Tagging the fish

For the smallest fish (Fig 4A, B), we placed the fish on a paper towel wet with aquarium water under a dissection scope. Using forceps to hold the minutien pin perpendicular to the fish, we pushed the pin into the dorsal muscle. With the pin still in place, we repositioned the fish to be upright and pulled the pin through the other side of the fish. Using forceps to hold the fishing line, we immediately pushed the line through the hole in the muscle made by the pin. If the tag was marked by knots or nail polish “beads,” we were careful not to pull those through the fish, causing more tissue damage. For larger fish (Fig 4C), where the needle can be attached to the fishing line, we followed a similar procedure. With the fish lying flat on a paper towel wet with aquarium water, we used forceps to hold the needle perpendicular to the fish and pierced the dorsal muscle. Shifting the fish to an upright position, we pushed the needle through the muscle (Fig 4D), pulling the needle and the line through the other side of the fish until 3-4 cm of line remained (Fig 4E). A sharp razor blade was used to trim the excess fishing line.

**Figure 4:**
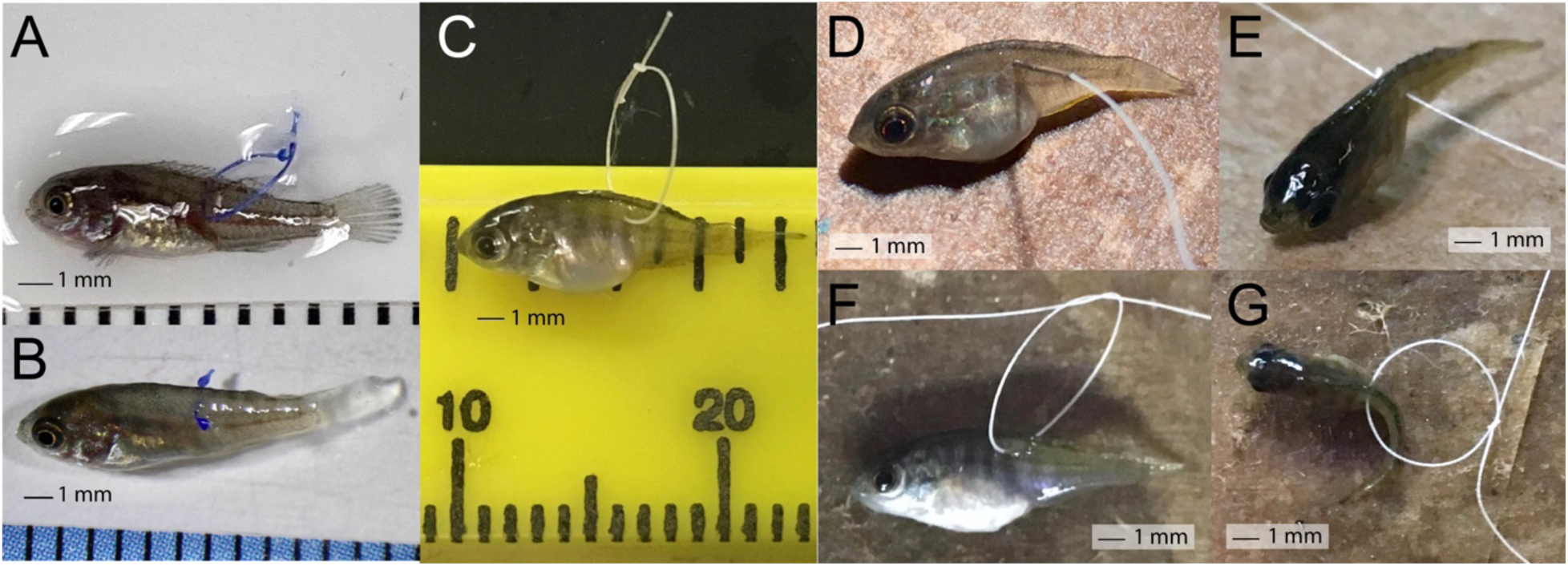
Tagging with a fishing line piercing. A) A juvenile tagged with a blue fishing line loop 1 hr after being removed from the mother’s buccal cavity (8.88 mm standard length, SL). B) A young juvenile (4 days post-removal from the mother’s buccal cavity) tagged with a blue fishing line barbell (10.08 mm SL). C) An older juvenile tagged with a fishing line loop (10.5 mm SL) on a ruler (inches top, mm bottom). Procedure for tagging larger juveniles: D) Pierce the dorsal muscle with the needle. E) Pull the needle and line through the dorsal muscle. F) Make a loop in the line and tie the first half of a square knot. Adjust the loop to the desired size. G) Finish the square knot. Cut the excess line using a razor blade. The same individual fish is pictured in panels C-G.

Once the fishing line was through the dorsal muscle of the fish, we tied it into a barbell (Fig 4B) or into a loop (Fig 4A,C) using a square knot (Fig 4F,G). If using a loop, we were sure to tie a large enough loop to allow the dorsal fin to raise and lower. However, a loop that was too large could get tangled in the tags of other fish or tank features (e.g., plastic plants). For the smallest fish, we found it safest to tie the barbell knot with two people. While one person pinched the line with forceps on the side of the fish where the second knot was to-be-tied, the second person tied a knot in the free line and slid the second knot into position using forceps. In general, if the knots were challenging to tie, we loosely pre-tied and then untied a knot in the line prior to threading it through the fish. For longer experiments, we placed a drop of super glue on the knot(s) to ensure they stayed tied, which dried in less than a minute. We kept the fish’s gills and body away from any glue and applied aquarium water to the gills, as needed, using a transfer pipette. As soon as possible, the fish was placed in a beaker of fresh aquarium water to recover. A fishing line tag can be removed quickly, and without anesthesia, by briefly removing the fish from its tank and cutting the line with a sharp razor blade.

#### Open field exploration

A physical tag may be more likely to affect locomotion and behavior, particularly in the smallest juveniles. We used an open field exploration to compare between juveniles that were anesthetized and tagged (n=7; one animal died between recovery and the behavioral test on the subsequent day) and those that were anesthetized but not tagged (n=8). This test is commonly used across species to assess movement, activity, and anxiety-like behavior (Prut and Belzung, 2003; Cachat et al., 2010; Solomon-Lane and Hofmann, 2019). We tagged the juveniles less than 2 hrs after being removed from the mother’s buccal cavity, and the open field test was carried out the next day (∼20 hrs later). The time it took to complete each tagging surgery was recorded (average 257 ± 22 s), and each control fish was kept out of water and manipulated with forceps for the same amount of time as one of the tagged fish. Additional MS-222 (or water) was applied to the fish during the procedure, as needed.

For the open field exploration, we placed fish individually in a small, novel aquarium (22.9 x 15.2 x 15.2 cm) without a cover for 30 min. For analysis, aquaria were divided into 4 zones: the territory zone, which contained a small terracotta pot territory / shelter, the close zone, the far zone, and the investigate zone (Fig 5A). Video cameras (Warrior 4.0, Security Camera Warehouse, Asheville, NC) recorded behavior from above, and we used BORIS (Friard and Gamba, 2016) to score the number of times fish entered each zone of the aquarium, and how long fish spent in each zone of the tank during a 10 min observation (minutes 20-30 of the test).

**Figure 5:**
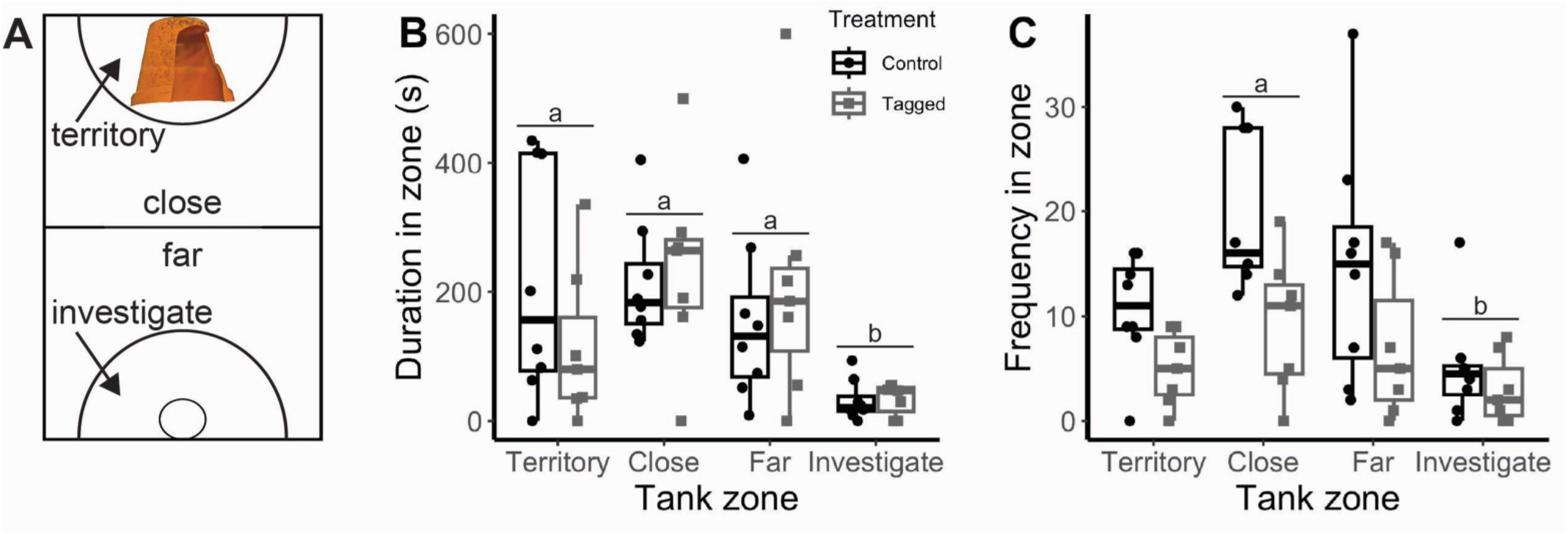
Open field exploration. **A)** Tank layout of the open field exploration, with the territory, close, far, and investigate zones. The territory zone has a small terracotta pot territory / shelter. **B)** Time spent (s) in each tank zone. **C)** Frequency entering each tank zone. Tagged fish (n=7) were anesthetized and tagged with a fishing line piercing less than 2 hrs after being removed from the mother’s buccal cavity. Control fish (n=8) were anesthetized but not tagged. The behavior test took place ∼20 hrs post-surgery. Different letters indicate significant differences.

### Statistics

All statistical analyses were conducted using R Studio (R version 4.4.0) (RStudio Team, 2022). Results were considered significant at the p<0.05 level, and averages ± standard error of the mean are included in the text. To compare anesthetization and recovery in control and tattooed juveniles, we used t-tests for data that met the assumptions of parametric statistics. A Wilcoxon signed ranks test was used for data that did not meet those assumptions. Cohen’s d was reported to estimate effect size for significant pairwise differences (small effect: 0.2 < d < 0.5; medium: 0.5 < d < 0.8; large: 0.8 < d). Pearson correlations were used to test for associations among recovery metrics in control and tattooed fish. To compare locomotion and behavior in the open field exploration between control and fishing line tagged juveniles, we used a two-way mixed ANOVAs, with tagging treatment as the independent measure and aquarium zone (frequency entering, time in) as the repeated measure. Tukey’s HSD test was used for *post hoc* analysis of significant ANOVA results. Eta-squared was reported to estimate effect size for significant ANOVA results (small effect: 0 < η^2^ < 0.01; moderate: 0.01 < η^2^ < 0.06; large: 0.06 < η^2^).

## RESULTS

### Recovery from anesthesia and injection of tattoo ink

To investigate the safety and efficacy of the tattoo ink injection procedure on very young, very small juveniles (< 2 weeks after removal from the mother’s buccal cavity), we compared the anesthetization and recovery of fish that were both anesthetized and tagged (tagged: n=8) to fish that were anesthetized but not injected (control: n=8). Fish in both treatment groups were removed from the MS-222 immediately after ventilation stopped and kept out of water for 1 min on a paper towel wet with aquarium water. Tagged fish were injected during the time out of water.

While being anesthetized, juveniles first lost equilibrium (30.6 ± 2.9 s), after which, body movement (52.5 ± 3.0 s) and gill ventilation (54.1 ± 3.5 s) stopped in close succession. Some individuals appeared to stop ventilating first, while others appeared to stop moving first. The treatment groups did not differ in time to lose equilibrium or time to stop moving (Table 1, Supplemental Fig 4A, B). Fish in the tagged group (although they had not been tagged yet) took significantly longer to stop ventilating (Supplemental Fig 4C), but overall, the groups did not differ in the total time spent in MS-222 (Table 1, Supplemental Fig 4D).

**Table 1:** Treatment differences in anesthetization and recovery between juveniles that were anesthetized (control) vs. juveniles that were anesthetized and tattooed. All juveniles remained out of the water for 1 min, whether or not they were tagged. Results of pairwise comparisons using t-tests (t) or a Mann-Whitney Wilcoxon test (W). Cohen’s d is reported for effect size of significant results. Significant results are in bold.

| Behavior | DF | Test Statistic | p-value | Effect size |
| --- | --- | --- | --- | --- |
| **Anesthetization in MS-222** |  |  |  |  |
| Loss of equilibrium | 10.963 | t = -0.35 | 0.73 |  |
| Stop moving | 9.50 | t = -1.07 | 0.31 |  |
| Stop ventilating | **9.01** | **t = -2.42** | **0.039** | **-1.3** |
| Total time in MS-222 | 10.07 | t = -1.388 | 0.20 |  |
| **Recovery** |  |  |  |  |
| Initial ventilation during artificial ventilation | 12.13 | t = 0.51 | 0.62 |  |
| Independent ventilation | 12.9 | t = 0.11 | 0.91 |  |
| Body movement | 13.47 | t = 0.90 | 0.38 |  |
| Initial regained equilibrium | 12.82 | t = 0.10 | 0.92 |  |
| Sustained regained equilibrium | 12.57 | t = -0.52 | 0.61 |  |
| Full recovery | 1 | W = 18 | 0.28 |  |

All of the fish in this cohort successfully recovered. After being placed in the recovery beaker, opercular movement during artificial ventilation occurred first (40.4 ± 9.5 s), followed by independent ventilation (without the aid of artificial ventilation, 107.9 ± 18 s). Initial body movement (128.4 ± 17.2 s), initial regained equilibrium (145.3 ± 21.9 s), and sustained regained equilibrium (162.6 ± 26.8 s) occurred next, in close succession. By definition, full recovery occurred last (407.3 ± 51 s). There were no significant differences for any recovery metric between fish that were anesthetized compared to those that were anesthetized and tagged (Table 1, Fig 3A, B).

Some recovery metrics were good predictors of later recovery (Table 2, Supplemental Fig 5). The timing of independent ventilation was significantly and positively associated with first body movement (Fig 3C) and initial regained equilibrium for both control and tagged fish, as well as sustained regained equilibrium in control fish. Initial body movement was significantly and positively associated with initial regained equilibrium in control and tagged fish, as well as sustained regained equilibrium in control fish. Initial regained equilibrium was significantly and positively associated with sustained regained equilibrium in control and tagged fish. Initial (Fig 3D) and sustained equilibrium were the only metrics associated with full recovery, and the associations were significant only for control fish. The timing of initial ventilation (while artificial ventilation was ongoing) did not predict any later recovery points (Table 2).

**Table 2:**
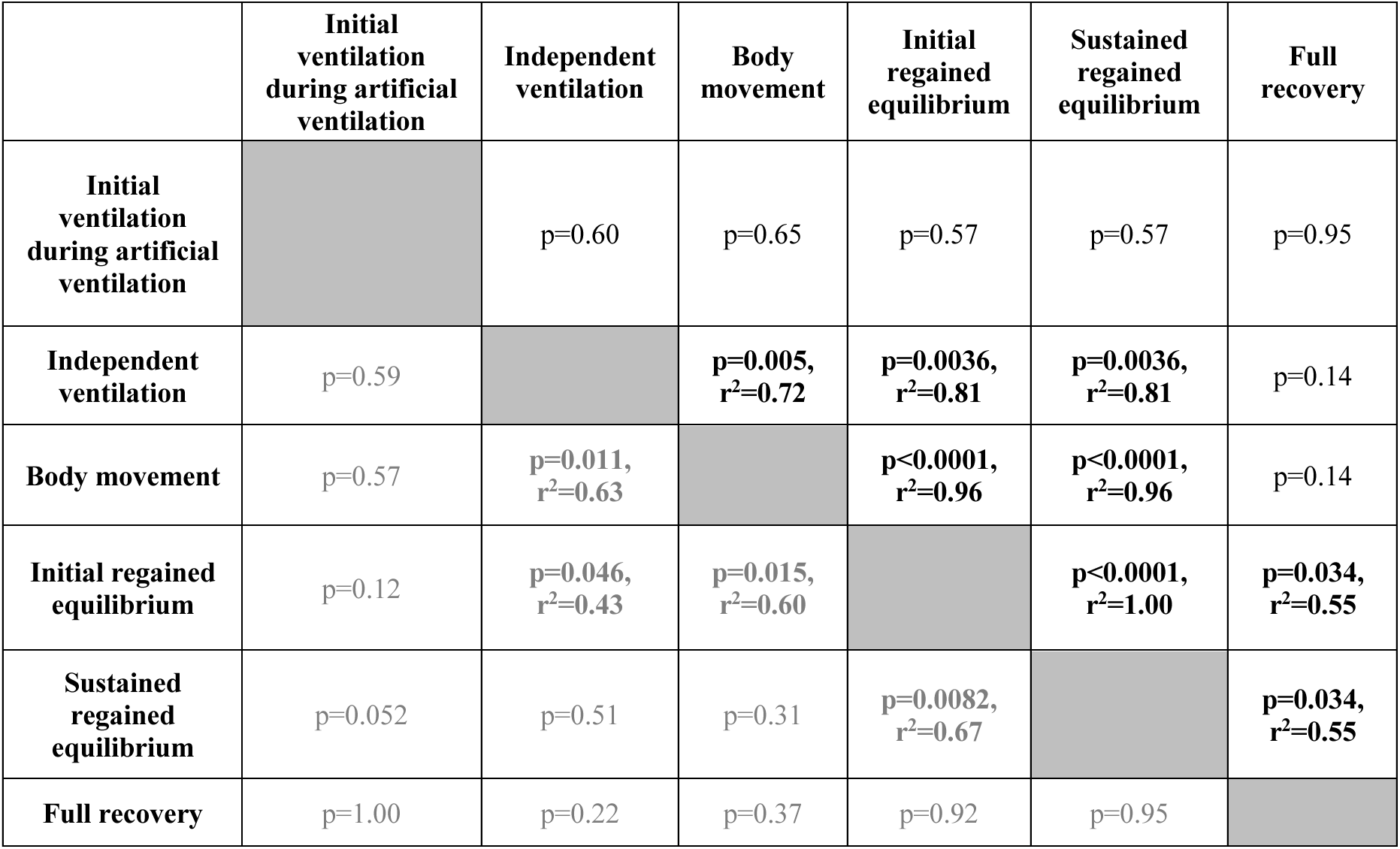
Associations between recovery metrics following anesthesia. Correlations for control fish in the upper triangular (black text). Correlations for tattooed fish in the lower triangular (grey text). Significant associations in bold. Adjusted r^2^ values shown for significant results.

### Open field exploration behavior with a fishing line piercing tag

We measured the time spent in each tank zone (Fig 5A), as well as the frequency entering each aquarium zone, for control and juveniles tagged with fishing line. We found a significant main effect of aquarium zone for time spent in each zone (F(1.46, 13.13)=13.44, p=0.001), but no main effect of treatment (F(1,9)=1.20, p=0.30), and no interaction effect (F(1.46, 13.12)=1.35, p=0.28). *Post hoc* analysis showed that juveniles spent significantly less time in the investigate zone compared to the territory (p=0.00026), close (p<0.0001), and far zones (p=0.00029). The other pairwise comparisons were not significant (close-territory: p=0.39, far-territory: p=1.00, far-close: 0.31) (Fig 5B). Similarly for the frequency of entering each zone, we found a significant main effect of aquarium zone (F(1.13, 10.2)=16.79, p=0.002), but no main effect of treatment (F(1,9)=1.90, p=0.20), and no interaction effect (F(1.13, 10.2)=0.48, p=0.53). Juveniles spent significantly less time in the investigate zone compared to the close zone (p=0.0004). The other pairwise comparisons were not significant (close-territory: p=0.23, far-territory: p=1.00, investigate-territory: p=0.088, far-close: p=0.21, investigate-far: p=0.082) (Fig 5C).

## DISCUSSION

Although there are a variety of approaches used to tag fish and other organisms, until the present study, no appropriate procedures were available for using these techniques to the youngest, and smallest, juveniles. We expect other researchers working with small organisms have encountered similar challenges. The procedures we describe here have allowed us to utilize two types of tags for the first time: an injected, subcutaneous tag (tattoo method) and a physical tag embedded in the body that remains externally visible (piercing method). These tags have made it possible for us to study the behavioral development and underlying neuromolecular mechanisms of juvenile *A. burtoni* (e.g., Solomon-Lane et al., 2022).

### Tattoo ink injection tagging

Injecting tattoo ink into the dorsal muscle of very young, small fish immediately creates a vibrant visual mark, allowing for individual identification. Initially, the tag was visible on both sides of the fish, but over time, the tag was visible for most fish just on one side of the body (e.g., Fig 2E). Tattoo ink is inexpensive and commercially available in an array of colors and mixing inks may be useful in creating more unique colors. Depending on the size of the fish, injection location may also be a useful unique identifier. We found the main limitation to be the fading of the colored mark over time, with lighter colors (e.g., yellow) appearing to fade more quickly than darker colors (e.g., black, blue). Our longest lasting tattoo tags persisted for 3-4 weeks (Fig 2E). Different colored inks, and different manufactures, have different compositions (Moseman et al., 2024), which could underlie the differences in tag visibility we observed. The different ink components could also have differential health effects (Grant et al., 2015; Moseman et al., 2024). In humans, tattoo ink is injected into the skin, specifically the papillary layer of the dermis. Over time, ink particles move deeper into the reticular dermis, and the tattoo persists nearly permanently because sufficient pigment stays in the skin (Grant et al., 2015; Moseman et al., 2024). Here, we injected ink into the dorsal muscle, which resulted in a moderately long-lasting tag, but it was not permanent. Pigment may disperse more readily, or fail to accumulate, in muscle compared to skin. Muscle can also change dramatically during fish development, including in muscle fiber type, size, and number (Patruno et al., 1998; Johnston, 1999). Once fish have sufficient dorsal muscle, a visible implant elastomer tag is a more permanent option than tattoo ink.

This procedure is fast and effective, and an experienced practitioner can tag a fish in less than 30 s. The use of anesthesia is necessary for the welfare and recovery of the juvenile fish, and we chose to use MS-222, a common and safe fish anesthetic (Wagner et al., 2003; Kiessling et al., 2009; Sneddon, 2012; Topic Popovic et al., 2012; Zahl et al., 2012) that can be cleared from circulation in less than 300 s (Kiessling et al., 2009). Observing individuals during anesthetization and recovery is highly informative for establishing an effective anesthetic dose, determining the optimal procedural timing, and ensuring full recovery (Sneddon, 2012). For example, some fish species will recover unaided from MS-222 (e.g., bluebanded gobies, *Lythrypnus dalli*, Solomon-Lane and Grober, 2012); however, *A. burtoni* juveniles recover best with artificial ventilation and continuous monitoring until full recovery. These data can also be used to identify and/or control for unintended biases. Here, we found that juveniles in the tagged group took longer to stop ventilating, even though they had not yet experienced anything different from the control fish (i.e., no fish had yet been tagged). With this information, we can test for downstream consequences. For example, despite this significant difference, tagged fish did not spend more total time in MS-222.

We found no significant differences in recovery timing between anesthetized control fish compared to anesthetized and tagged fish. This suggests that, overall, tattoo ink injections are safe, with limited negative physiological consequences. We did identify some effects of tagging on the temporal progression of recovery. In control fish, for example, the timing of initial and sustained regained equilibrium was significantly associated with the timing of full recovery. In contrast, for tagged fish, no recovery metric predicted the timing of full recovery. These differences support our approach of using recovery data as an indicator of how an individual was impacted by the procedure and if it is ready to return to its home aquarium or engage in an experimental test (Solomon-Lane and Grober, 2012). For example, we found that the timing of first opercular movement (initial ventilation) during artificial ventilation was not a predictor of subsequent stages of recovery. In contrast, independent ventilation and body movement, which occurred in close succession, predicted when equilibrium was regained. Additional validations of this procedure may be needed, depending on the species or experiment. For example, in our future studies related to social behavior, we will test whether tag color affects how fish behave or are treated by other members of their social group.

In addition to tattoo ink, we also tested the efficacy of injecting acrylic paint (Lotrich and Meredith, 1974), thinned slightly with Liquidtex® Flow Aid^TM^ to prevent the paint from drying rapidly in the needle. Acrylic paint tags lasted approximately two weeks, with the longest tag lasting four weeks, under our conditions. Acrylic paint is non-toxic, inexpensive, and available in many colors. Injections of Alcian blue (Thermo Scientific Chemicals) lasted approximately three weeks. With no color options other than blue, this did not meet our needs for distinguishing among multiple juveniles in a social group. Injections of food coloring lasted only a couple of days (data not shown).

### Fishing line piercing tagging

Piercing the dorsal muscle with a fishing line tag is a highly effective way of individually marking juvenile fish that can last for months. The materials are inexpensive and commercially available, there are multiple methods for creating unique tags (e.g., line color, “bead” number, placement, and/or color(s)), and the surgery is quick, taking an experienced researcher less than 2 min to anesthetize and tag a fish. Recovery and survival is extremely high in older juveniles (e.g., Solomon-Lane et al., 2022), and although very young, small juveniles are more fragile, recovery and survival were also high in this cohort. As needles and fishing line (or similar, e.g., 44-gauge stainless steel wire, Remington Industries) continue to get thinner and lighter, we expect this method can become even less invasive for very small fish. For a short experiment (< 1 week), line colored with permanent marker remained vibrant under our aquarium conditions, but the colors did fade after 3-4 weeks. When super glue was added over a knot, the color under the glue remained visible for much longer (at least 8 weeks).

A physical tag may be more likely to affect locomotion and behavior than a fully internal tag (e.g., tattoo ink); therefore, we tested fish in an open field exploration under conditions that we considered most likely to reveal potential negative effects on behavior. Juveniles were tagged less than 2 hrs after being removed from the mother’s buccal cavity, and at the time of the open field test, they had less than 24 hrs experience as freely-swimming fish. Tagging did not have a significant effect on where juveniles spent time in the aquarium, or how frequently they entered the different aquarium zones. Interestingly, tagged juveniles entered the territory zone and close zone fewer times than control fish. Remaining in or near the territory can be associated with shyness (Harmon et al., 2024), and injured fish may be more likely to hide in the territory.

However, the effect we saw was in the opposite direction. This behavior pattern could indicate that locomotion decreases for a period of time after tagging, therefore, we plan to measure locomotion as a control in future studies. Depending on the experiment, species, and developmental stage, a longer (or shorter) recovery period may be needed, and testing relevant behaviors on subsequent days post-surgery will be valuable for determining appropriate experimental timing. Overall, we were impressed by the quick recovery of these very small, very young juveniles.

## Conclusion

Overall, the lack of appropriate procedures for individually identifying juveniles this small and young likely has contributed to the relative paucity of early-life studies, in this and other small fish species. In social neuroscience, behavioral ecology, animal behavior, developmental biology, fisheries research, and related fields, experiments that follow individuals through development are critical to uncovering the emergence of individual phenotypic variation, as well as the underlying mechanisms and fitness consequences (Taborsky, 2016). In describing these procedures, we hope they will be useful to other researchers and will continue to be improved in the future.

## ACKNOWLEDGEMENTS

We thank Isaac Miller-Crews, Gabriel Rocha, Cameron Hamson, and Caitlyn Kwun for size measurements of juvenile *A. burtoni*. We thank Dr. Sarah Budischak for use of her camera rig and microscope for images of tattoo ink injected fish. We thank members of the Solomon-Lane Lab for assisting with artificial ventilation during the development of these methods, fish care, and pictures of tagged fish, including Veronica Britton, Alex Kasiske, Jessica Gonzalez Rodriguez, Eliyah Stern, and Andrew Yuan. Special thanks to Dr. Mel Coleman for methodological suggestions. We also thank members of the Hofmann Lab and Solomon-Lane Lab for contributing to animal care and maintenance. This work was supported by NSF grant IOS-1354942 to HAH, the BEACON Center for the Study of Evolution in Action awards #947 (2016) and #1081 (2017, 2018) to TKSL and HAH, NSF grant IOS-2341006 to TKSL,

Department of Natural Sciences funding to TKSL, and the Keck Summer Research Fellowship to DDB.

## ETHICS

All protocols and procedures employed in this study adhered to the ARRIVE guidelines for reporting study design and statistical analysis; experimental procedures; experimental animals and housing and husbandry (Percie du Sert et al., 2020) and in compliance with the US National Research Council’s Guide for the Care and Use of Laboratory Animals, the US Public Health Service’s Policy on Humane Care and Use of Laboratory Animals, and Guide for the Care and Use of Laboratory Animals. All procedures were ethically reviewed and approved by the Institutional Animal Care and Use Committee (IACUC) of the Department of Natural Sciences of Pitzer and Scripps Colleges (protocol #19-001) and the IACUC of The University of Texas at Austin (protocol # AUP-2015-00153).

## DATA AVAILABILITY

All relevant data are available for reviewers at http://datadryad.org/share/siBW2kDGrSr-wVZ-hUoRsYuk-sL55N0ErGK89YRoKjQ. These data will be made public upon acceptance for publication.

