## Supplemental Information for "Tagging Very Small Fish: Two Effective and Low Impact Methods"

**Supplemental Information for: Tagging Methods for Very Small Fish**

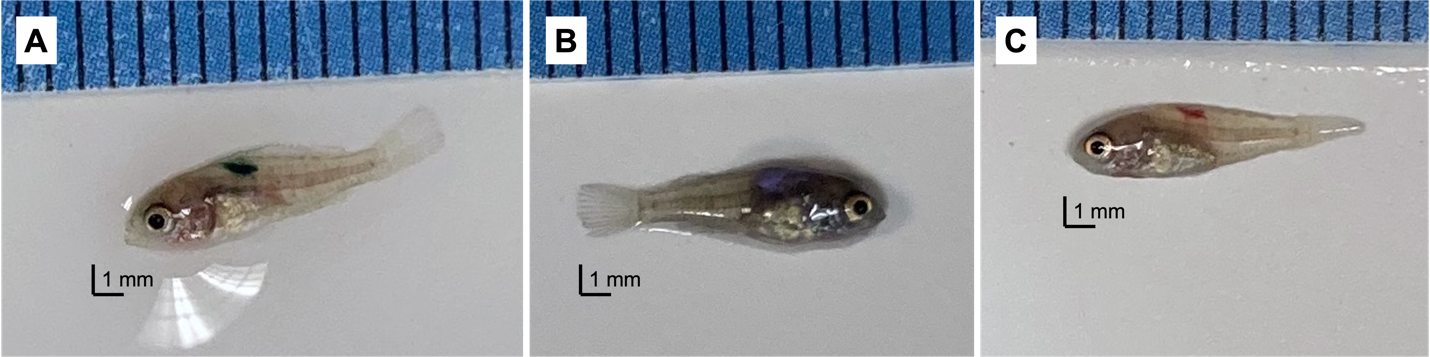

**Supplemental Figure 1:** Example images of juvenile fish (at the same scale) 1 day after tattoo ink injection tagging with **A)** green ink, **B)** purple ink, and **C)** red ink.

**Detailed set-up information for the Nanoject II Auto-Nanoliter Injector**

1) Attach a magnetic base to the bottom of the micromanipulator, place on the magnetic base plate, and turn on the switch to secure the micromanipulator in place (Fig 2A, Supplemental Fig 2E).

2) Secure the Nanoject in the micromanipulator.

3) Set the Nanoject settings to 50.6 nL, “fast” injection using the switches on the side of the controller.

4) Prepare the needle:

a) Unscrew the black tip of the Nanoject and use a dental tool to remove the 3 O rings. The top O ring is flexible, black, and thin. The middle O ring is hard, white plastic. The third O ring is flexible, black, and thicker than the top one (Supplementary Figure 2D).
b) Using forceps under a dissecting scope, break the tip off of a capillary tube needle.
c) Backfill the capillary tube needle with mineral oil using a ~26-28 gauge needle. Fill the whole capillary tube. Avoid bubbles by continuing to inject mineral oil while slowly withdrawing the needle from the capillary tube.
d) Put the back end (the flat, non-needle side) of the capillary tube needle through the black tip of the Nanoject and replace the O rings. Slide the thick black O-ring down the capillary tube needle (~1 cm). The white O-ring has a small indentation on one side that fits around the capillary tube. The thin black O-ring sits on top of the white O-ring.
e) Thread the metal plunger on the Nanoject through the holes in the O rings and into the capillary tube needle. Screw the black tip into place. Do not force it, and make sure the threads are lined up correctly.
f) Press “empty” on the Nanoject controller. This will cause the mental plunger to depress, dispelling mineral oil from the needle. Depress the plunger about 50%-70% of the way.
g) Wipe the needle clean with a Kim wipe and ethanol. Let the ethanol dry briefly.
h) Prepare the tattoo ink on Parafilm or in a weigh boat right before the injection process begins to avoid it drying out. The commercially available tattoo inks have different consistencies and drying times. For example, we found the Millennium Mom’s green and yellow tattoo inks to have a slower drying time in the needle compared to red and blue. Green tattoo ink was noted to be the most vibrant and easily identifiable color. The blue ink tended to last the longest. For *Astatotilapia burtoni*, or another neutrally colored fish, the neutral colors (e.g., grey, white) were not very visible and are not recommended. Dilute the tattoo ink with water or Keep it Wet, if needed, and mix well.
i) Put the tip of the needle in the tattoo ink and press “fill” on the Nanoject controller. Do not load more of the ink than needed for 4-8 injections.
j) Perform a test injection by pressing “inject” on the Nanoject controller 1-3 times to make sure everything is working. The most common complication is a clogged needle or ink that is too thick to eject.
k) Clean the needle with ethanol using a Kim wipe before injecting a fish.

5) Move the microscope into place next to the Nanoject and position the lights. Make sure you can see the tip of the needle in the scope. Move the manipulator up and down to make sure that when you inject, the tip of the needle and the animal’s dorsal muscle are still in view.

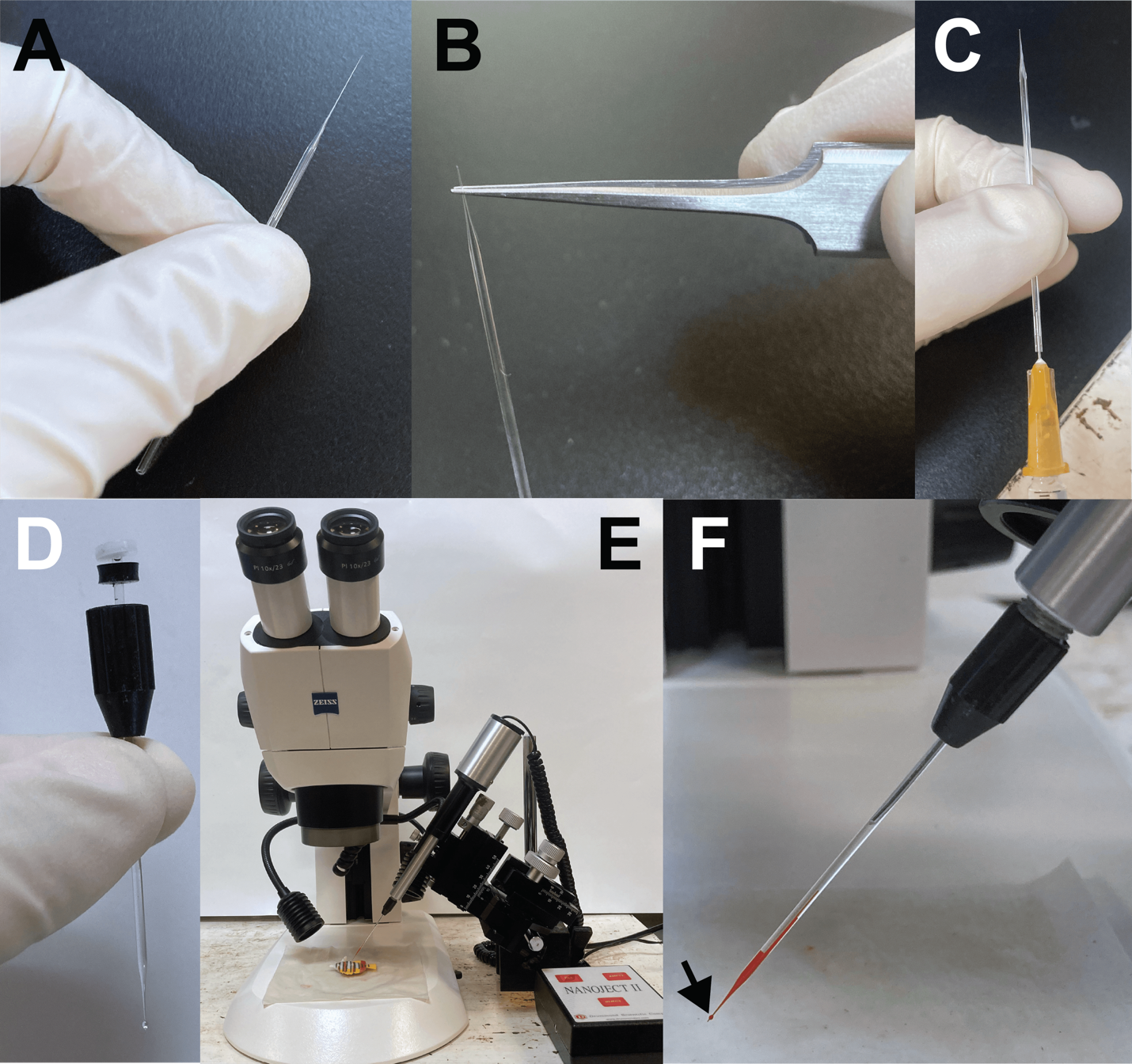

**Supplemental Figure 2:** **Prepare needle prior to injection.** A) Pulled capillary tube needle. B) Using forceps, break the tip off a capillary tube needle. C) Backfill the capillary tube needle with mineral oil using a ~26-28 gauge needle. D) Place the needle inside of the black tip of the Nanoject and then replace the O rings, in order, around the capillary tube needle. E) Nanoject injector set up. F) Prepared needle with mineral oil at the top and loaded with tattoo ink.

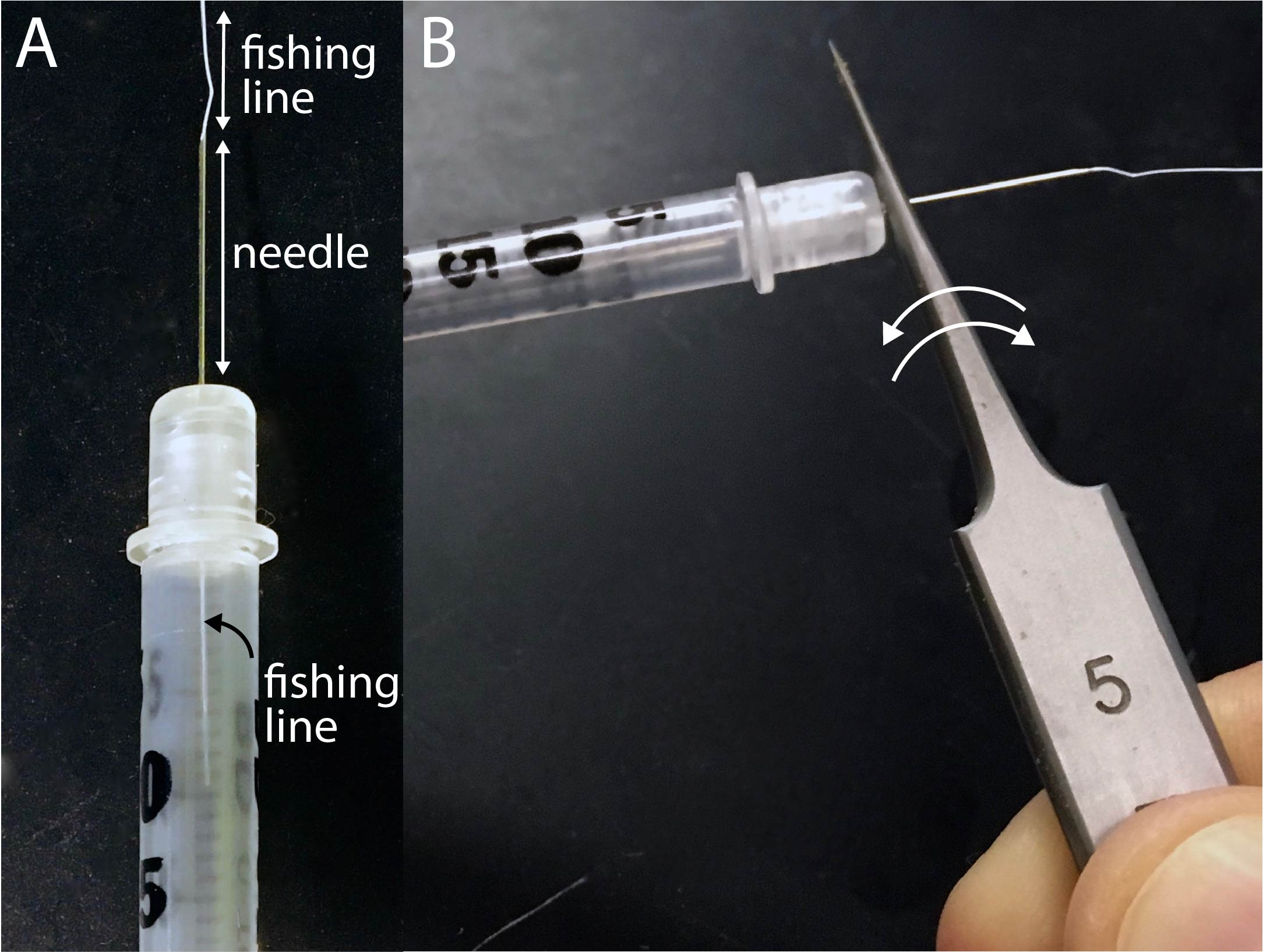

**Supplemental Figure 3: Making the fishing line tag.** A) Thread the fishing line through the needle of an insulin syringe until it is visible in the barrel of the syringe. B) Use forceps to grasp the base of the needle. Rock back and forth (arrows) until the needle breaks off from the syringe. Not shown: From the blunt side of the needle, pull the line through, until it no longer sticks out beyond the needle tip. Use a razor blade to cut off excess line.

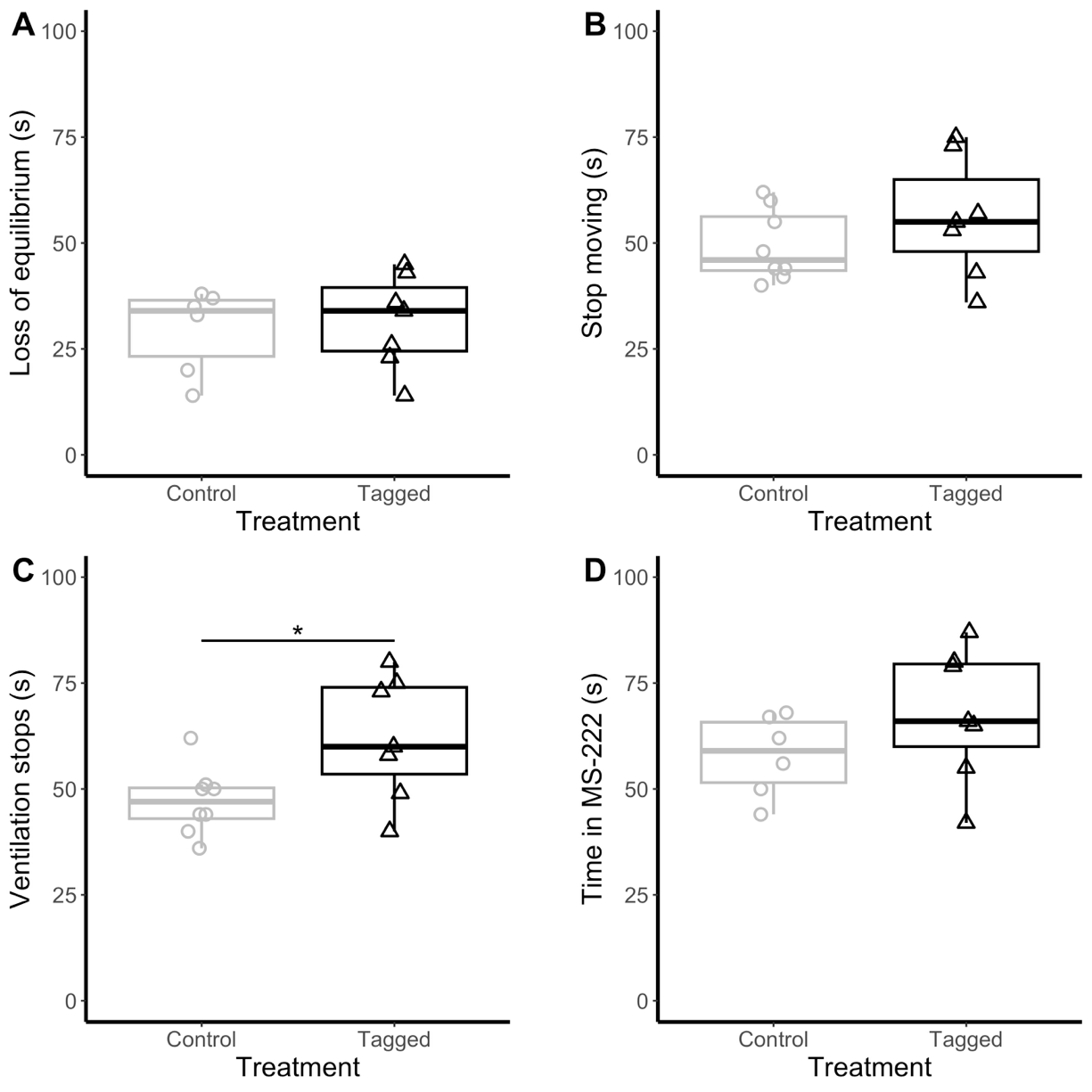

**Supplemental Figure 4:** Time in MS-222 until A) loss of equilibrium (t=-0.35, df=10.96, p=0.73), B) body movement stops (t=-1.07, df=9.50, p=0.31), and C) ventilation stops (t=-2.42, df=9.01, p=0.039). D) Total time in MS-222 (t=-1.39, df=10.07, p=0.20). *p<0.05

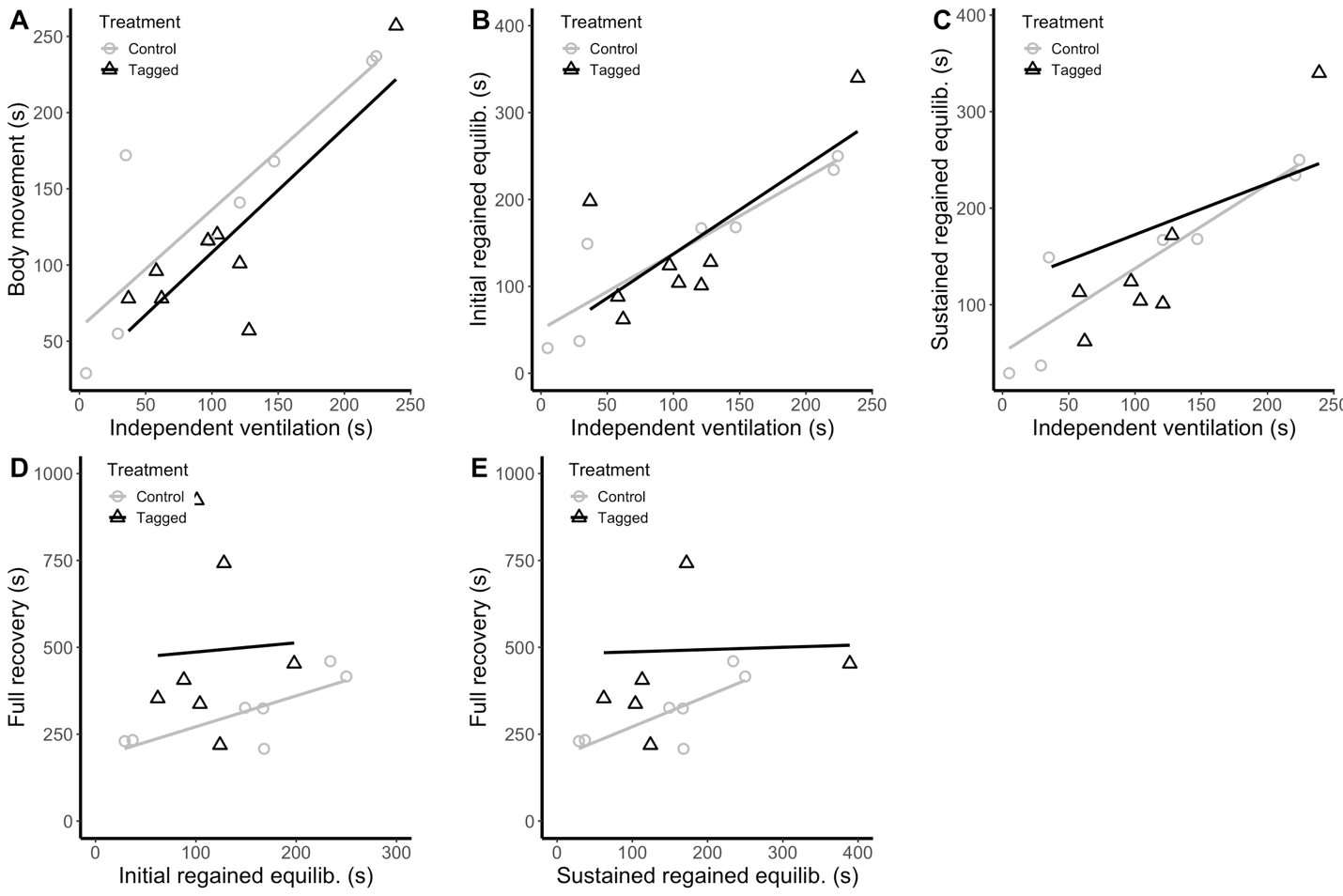

**Supplemental Figure 5:** Associations among the timing of recovery stages following anesthesia: A) independent ventilation and initial body movement, B) independent ventilation and initial regained equilibrium, C) independent ventilation and sustained regained equilibrium, D) initial regained equilibrium and full recovery, and E) sustained regained equilibrium and full recovery.
